# Mutations in *TAC1B* drive *CDR1* and *MDR1* expression and azole resistance in *C. auris*

**DOI:** 10.1101/2025.02.11.637698

**Authors:** Katherine S. Barker, Darian J. Santana, Qing Zhang, Tracy L. Peters, Jeffrey M. Rybak, Joachim Morschhäuser, Christina A. Cuomo, P. David Rogers

## Abstract

**Objective:** *Candida auris* has emerged as a fungal pathogen of particular concern owing in part to its propensity to exhibit antifungal resistance, especially to the commonly prescribed antifungal fluconazole. In this work we aimed to determine how mutations in the transcription factor gene *TAC1B*, which are common among resistant isolates and confer fluconazole resistance, exert this effect.

**Methods:** Selected *TAC1B* mutations from clinical isolates were introduced into a susceptible isolate and reverted to the wild-type sequence in select clinical isolates using CRISPR Cas9 gene editing. Disruption mutants were likewise generated for select genes of interest. *TAC1B* mutants were subjected to transcriptional profiling by RNA-seq, and relative expression of specific genes of interest was determined by qRT-PCR. Antifungal susceptibilities were determined by modified CLSI broth microdilution.

**Results:** *TAC1B* mutations leading to A640V, A657V, and F862_N866del conferred fluconazole resistance, as well as increased resistance to other triazoles, when introduced into a susceptible isolate. RNA-seq revealed that the ATP-Binding Cassette (ABC) transporter gene *CDR1* as well as the Major Facilitator Superfamily (MFS) transporter gene *MDR1* were both upregulated by these *TAC1B* mutations. Disruption of *CDR1* greatly abrogated resistance in strains with *TAC1B* mutations whereas disruption of *MDR1* had little to no effect. However, disruption of both *CDR1* and *MDR1* resulted in an additional reduction in resistance as compared to disruption of either gene alone.

**Conclusion:** *TAC1B* mutations leading to A640V, A657V, and F862_N866del all result in increased resistance to fluconazole and other triazole antifungals, and increased expression of both *CDR1* and *MDR1* in *C. auris*. *CDR1* is the primary driver of resistance conferred by these *TAC1B* mutations.

## Introduction

Since its initial identification in 2009, *Candida auris* has emerged as an important healthcare-associated fungal pathogen causing outbreaks worldwide(Lockhart, Chowdhary, & Gold, 2023; Lyman et al., 2023). Of particular concern is the prevalence of antifungal resistance among *C. auris* isolates, including resistance to two and sometimes all three antifungal classes currently available for treatment of serious *Candida* infections(Ostrowsky et al., 2020; Rybak, Cuomo, & Rogers, 2022). Strikingly, over 90% of isolates exhibit resistance to the most widely prescribed antifungal worldwide, the triazole antifungal fluconazole.

Fluconazole exerts its antifungal activity by competitively binding to and inhibiting sterol demethylase, a key enzyme of the fungal sterol biosynthesis pathway. In *Candida* species, this results in both reduced production of the major membrane sterol ergosterol as well as accumulation of a toxic sterol intermediate(Kelly, Lamb, Corran, Baldwin, & Kelly, 1995). Fluconazole resistance in *Candida* species can be the result of mutations in the *ERG11* gene which encodes sterol demethylase, resulting in altered drug binding or enhanced preference for the natural substrate, leading to reduced enzyme inhibition(Marichal et al., 1999). Resistance can also be due to mutations in the genes encoding the transcriptional regulators Tac1, Mrr1, or Upc2 which result in overexpression of genes encoding the ATP Binding Cassette (ABC) transporter Cdr1, the Major Facilitator Superfamily (MFS) transporter Mdr1, or Erg11, respectively(Coste, Karababa, Ischer, Bille, & Sanglard, 2004; Dunkel et al., 2008; Morschhäuser et al., 2007). Rarely, loss-of-function mutations are found in the sterol desaturase gene *ERG3* conferring resistance by abrogating the need for sterol demethylase(Kelly, Lamb, & Kelly, 1997).

Three predominant *ERG11* mutations, leading to amino acid substitutions Y132F, K143R, and VF125AL, have been shown to contribute to fluconazole resistance in *C. auris* (Rybak et al., 2021). The transcription factor Mrr1 and the MFS transporter Mdr1 have also been implicated in resistance in isolates from Clade III(Li, Coste, Bachmann, Sanglard, & Lamoth, 2022). We have previously shown that *CDR1* deletion in a resistant isolate that overexpresses this gene leads to a significant reduction in resistance to triazole antifungals(Rybak et al., 2019). We subsequently identified mutations in the gene encoding the transcription factor Tac1B in resistant isolates that were evolved *in vitro* in the presence of fluconazole, observed similar mutations among resistant clinical isolates, and demonstrated that the mutation leading to the A640V substitution in and of itself confers increased fluconazole resistance(Rybak et al., 2020). However, the relationship between mutations in *TAC1B*, *CDR1*, or other potential resistance effectors remains unclear as the effect of *TAC1B* mutations on the expression of *CDR1* or other genes has yet to be investigated. In the present study we establish that the *TAC1B* mutations leading to A640V, A657V, and F862_N866del all result in increased resistance to triazole antifungals, as well as increased expression of both *CDR1* and *MDR1* in *C. auris*. We also show that *CDR1* is the primary driver of resistance conferred by these *TAC1B* mutations.

## Methods

### Isolates, strains, and growth conditions

The clinical isolates and strains described in this study are listed in **Supplementary Table S1**. Cells were propagated in YPD (1% yeast extract, 2% peptone, 2% dextrose) at 35°C and stored in 40% glycerol at -80°C.

**Strain construction.** *C. auris* strain construction was based on the methods described previously(Carolus et al., 2024; Lombardi, Oliveira-Pacheco, & Butler, 2019), and detailed methods are described in **Supplementary Methods**. Oligonucleotides used in this study are listed in **Supplementary Table S2**.

### Minimum inhibitory concentration (MIC) determinations by broth microdilution

MICs for fluconazole, itraconazole, voriconazole, posaconazole, and isavuconazole were measured by modified CLSI broth microdilution assays as described previously(Rybak et al., 2021).

### RNA isolation

An aliquot of cells from an overnight culture was used to inoculate 10 ml MOPS-buffered RPMI + 2% glucose (pH 7.0) to a OD_600_=0.08-0.12, followed by incubation at 35°C in a shaking incubator (220 rpm) until mid-log phase (6 hrs). Cell cultures were grown in triplicate. Cells were collected by centrifugation, supernatants removed, and cell pellets stored at -80°C. RNA was extracted using methods described previously with some modification(Santana & O’Meara, 2021). Cell pellets were resuspended in 100 µl FE Buffer (98% formamide, 0.01 M EDTA) at room temperature. Fifty microliters of 1 mm RNase-free glass beads were added to the cells and were subjected to vortex disruption for 5 min at room temperature followed by snap-cooling on ice. The cell lysate was clarified by centrifugation, and the supernatant was DNase-treated for 30 min followed by isopropanol precipitation. RNA integrity was confirmed by agarose gel electrophoresis, and concentrations were approximated by Nanodrop.

### cDNA synthesis and qRT-PCR

cDNA was synthesized from 500 ng RNA using the RevertAid First Strand cDNA Synthesis Kit (Invitrogen/Thermo Fisher) with the provided random primer mix according to the manufacturer’s instructions. qRT-PCR was performed from three biological replicates, each with three technical replicates, using SYBR Green PCR master mix (Bio-Rad). Fold changes were calculated using the ΔΔ^CT^ method with target gene CT values normalized to *ACT1* CT values (generating dCT values) and the median 1c dCT value (from the three biological replicates) for each target gene used as the comparator for fold change calculation.

### RNA sequencing and analysis

RNA sequencing was performed using Illumina NextSeq for stranded mRNA. Libraries were prepared with paired-end adapters using Illumina chemistries per manufacturer’s instructions, with read lengths of approximately 150 bp with at least 50 million raw reads per sample. Data were analyzed using the Galaxy web platform public server at usegalaxy.org*(Afgan et al., 2018)*. Read quality was assessed and reads were quality-trimmed with a PHRED cutoff of 20 using FastP(Chen, Zhou, Chen, & Gu, 2018). Reads were then mapped to the *C. auris* B8441 reference assembly (NCBI GCA_002759435.2) using RNA Star with default parameters (Dobin et al., 2013) followed by quantification using featureCounts (Liao, Smyth, & Shi, 2014) and assessment of differential expression using DESeq2(Love, Huber, & Anders, 2014). Expression fold-change and significance cutoffs of >2-fold up- or down-regulated and an adjusted p-value less than 0.05 were established to categorize dysregulated genes. Gene Ontology annotations were retrieved from the *Candida* Genome Database (Skrzypek et al., 2017) to identify significantly dysregulated GO terms using GoSeq(Young, Wakefield, Smyth, & Oshlack, 2010), normalizing for feature lengths extracted from the featureCounts output. Transcriptional overlap between strains was assessed using either UpsetR R Package (Conway, Lex, & Gehlenborg, 2017) or DiVenn 2.0(Sun et al., 2019). The GEO accession number was assigned as GSE288372.

## Results

### *TAC1B* mutations observed in clinical isolates confer fluconazole resistance and reduced susceptibility to other triazole antifungals

Previously we showed that introduction of the *TAC1B* mutation leading to the A640V substitution into susceptible clinical isolate AR0387 results in an increase in fluconazole MIC from 1 µg/mL to 8 µg/mL, and correction of this mutation to the wild-type sequence in resistant clinical isolate AR0390 reduces fluconazole MIC from 256 µg/mL to 16 µg/mL(Rybak et al., 2020). In order to further investigate the contribution of *TAC1B* mutations to triazole antifungal resistance, we introduced A640V, A657V, and F862_N866del into *C. auris* strain 1c. Strain 1c is derived from highly fluconazole-resistant Clade Ic clinical isolate Kw2999 (fluconazole MIC = 256 µg/mL) that harbors a *ERG11* mutation leading to the K143R substitution and the *TAC1B* mutation leading to the A640V substitution(Ahmad, Khan, Al-Sweih, Alfouzan, & Joseph, 2020). In strain 1c, both mutations have been corrected to their wild-type sequences resulting in fluconazole MIC of 2 µg/mL. As previously demonstrated in isolate AR0387, reintroduction of the A640V substitution in 1c (strain 1cA640V) resulted in an increase in fluconazole MIC to 32 µg/mL. Introduction of the A657V substitution (strain 1cA657V) likewise increased the MIC to 32 µg/mL, whereas F862_N866del (strain 1cADdel) increased the MIC to 64 µg/mL **(Table 1)**. The susceptibilities to voriconazole, isavuconazole, itraconazole, and posaconazole were also reduced **(Table 1)**. These results indicate that each of these *TAC1B* mutations confer increased resistance to the triazole antifungals.

**Table 1.** Fluconazole MIC values^1^ of Parental Strain 1c and its *TAC1B* Mutant Derivative Strains.

|  | Fluconazole | Itraconazole | Posaconazole | Voriconazole | Isavuconazole |
| --- | --- | --- | --- | --- | --- |
| 1c | 2 | 0.031 | 0.0625-0.125 | 0.015 | 0.0038-0.0078 |
| 1cA640V |  |  |  |  |  |
| Clone 1 | 32 | 0.25-0.5 | 0.125-0.25 | 0.25 | 0.125-0.25 |
| Clone 2 | 32 | 0.5 | 0.25 | 0.25 | 0.25 |
| 1cA657V |  |  |  |  |  |
| Clone 1 | 32 | 0.25 | 0.125 | 0.25 | 0.125 |
| Clone 2 | 32 | 0.5 | 0.125-0.25 | 0.25-1 | 0.125-0.25 |
| 1cADdel |  |  |  |  |  |
| Clone 1 | 64 | 0.25-0.5 | 0.125-0.25 | 0.5 | 0.5 |
| Clone 2 | 64 | 0.5 | 0.125-0.25 | 0.5 | 0.5 |

<sup>1</sup> MIC values are in µg/mL and represent at least three biological replicates

### Mutations in *TAC1B* drive overexpression of *CDR1* and *MDR1*

As there were similar shifts in azole resistance for each mutant, we investigated whether there was a common transcriptional response. We performed RNA-seq on strain 1c and clone A from each *TAC1B* mutant strain and established a cutoff for significant gene dysregulation at 2-fold compared to strain 1c with a DeSeq2 adjusted p-value of <0.05. We observed an overlapping set of dysregulated genes among the *TAC1B* mutant backgrounds, but each also showed substantial differences in transcriptional response. For instance, the A640V mutation resulted in only 5 dysregulated genes, each of which was also represented by either A657V or F862_N866del (ADdel) **(Figure 1A)**. Of the 40 ADdel mutant dysregulated genes, 72.5% were dysregulated in at least one of the other two strains **(Figure 1A)**. Interestingly, the A657V mutation resulted in a large unique transcriptional response, with 358 genes being dysregulated only in this mutant **(Figure 1A)**. Gene Ontology Enrichment analysis revealed similar patterns of enriched gene sets between strains, with common terms surrounding transmembrane transport and xenobiotic detoxification frequently represented, in line with the role of *TAC1* homologs in other species in regulating transporter and efflux activity **(Figure 1B)**. Among these were efflux pumps *MDR1*, upregulated in all three mutants, and *CDR1*, upregulated in the A640V and ADdel mutants. In the A657V mutant, *CDR1* showed an 1.7-fold increase in expression. While this didn’t meet the fold-change cutoff for significance, this increase may still be sufficient to functionally explain the observed increase in triazole MIC **(Figure 1B)**. Dozens of other genes with predicted transporter function were also dysregulated in this strain background, raising the possibility of other influential effectors **(Figure 1C)**.

**Figure 1.**
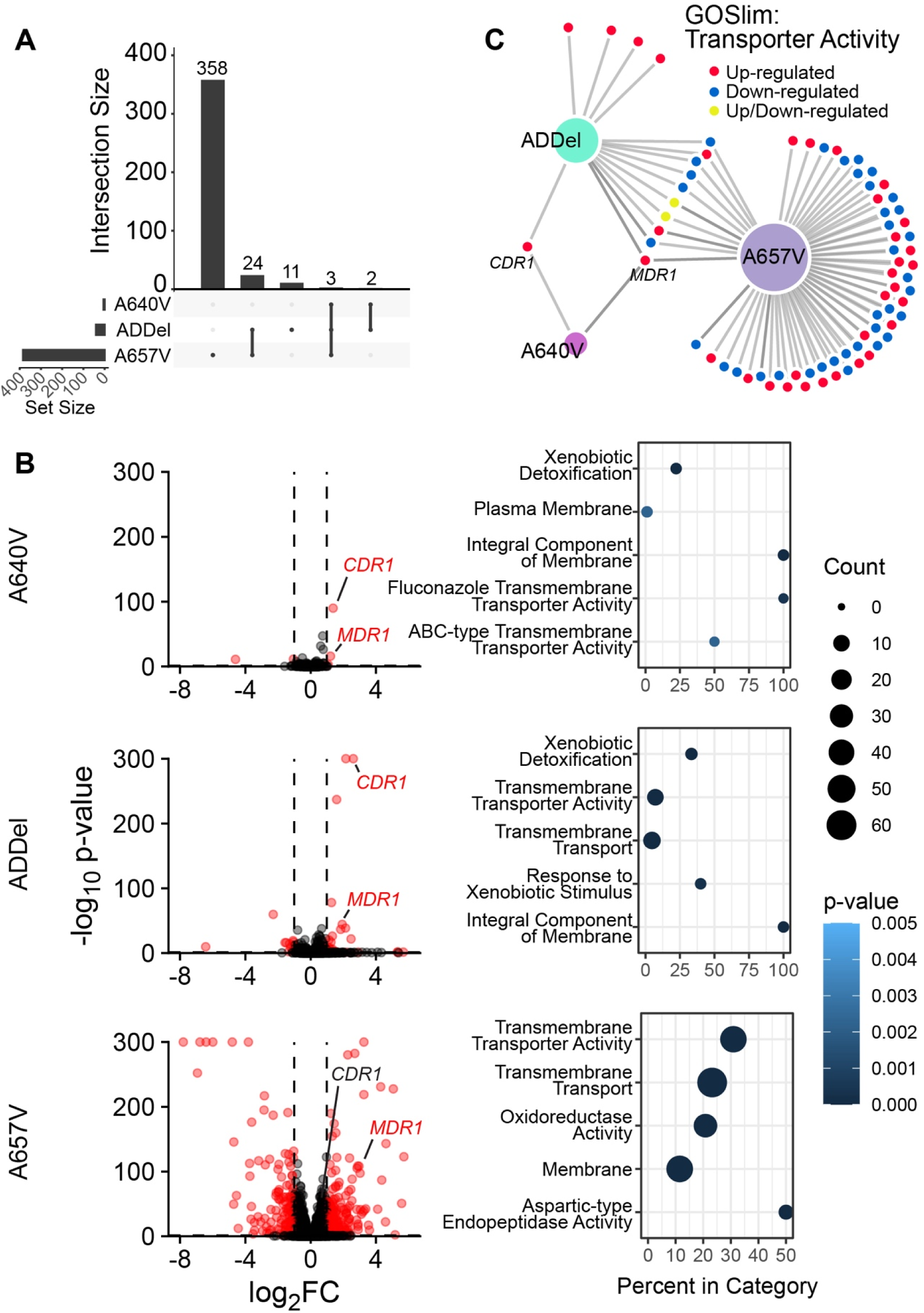
(A) UpSet plot of transcriptional network overlap, demonstrating the intersections of genes that are dysregulated by >2 fold and adjusted p-value of <0.05 compared to the parental isolate for each strain harboring different *TAC1B* mutations. Each bar in the upper region signifies the number of dysregulated genes in each intersection while the lower region signifies which strains exhibit dysregulation in those genes. **(B)** Volcano plots demonstrate magnitude of gene dysregulation for each strain, with significantly dysregulated genes colored in red. The top 5 overrepresented Gene Ontology terms in each dataset are listed. **(C)** All dysregulated genes that match the GOSlim term for Transporter Activity are represented for each strain. Colors represent the directionality of dysregulation. For all panels, strains are indicated by their harbored *TAC1B* mutation, while “ADdel” refers to the F862_N866del mutation.

We then measured *CDR1* and *MDR1* expression by qRT-PCR in strains 1cA640V, 1c657V, and 1cADdel compared to strain 1c and observed 2.5-, 1.6-, and 6-fold increase in *CDR1* expression (**Figure 2A**) and 2.6-, 11.7-, and 3.5-fold increase in *MDR1* expression, respectively (**Figure 2B**). We then measured *CDR1* and *MDR1* expression by qRT-PCR in susceptible clinical isolates AR0387, its derivative carrying the A640V substitution, resistant clinical isolate AR0390, and its derivative where the A640V mutation has been corrected to the wild-type sequence. *CDR1* and *MDR1* were upregulated 3.2-fold and 3-fold, respectively, in the presence of the A640V substitution in the AR0387 background (**Figure 2C and 2D**). Likewise, expression of these genes was reduced to wild-type levels when the A640V substitution was corrected to the wild-type sequence (**Figure 2C and 2D**). These data indicate that *TAC1B* mutations drive *CDR1* and *MDR1* overexpression in *C. auris* resistant isolates.

**Figure 2.**
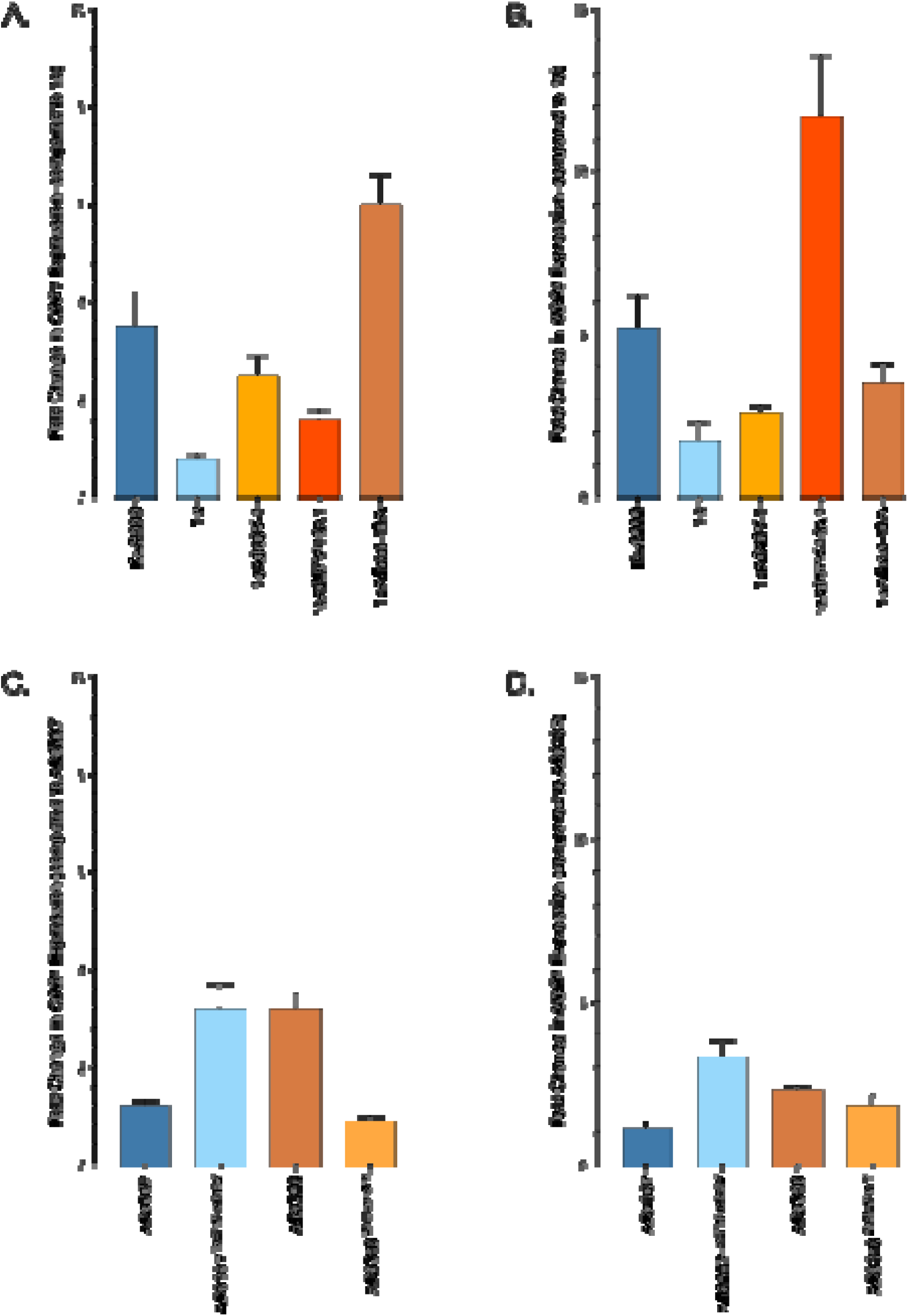
Fold change in *CDR1* **(A)** and *MDR1* **(B)** RNA expression for 1c *TAC1B* mutant strains and *CDR1* **(C)** and *MDR1* **(D)** transcript abundance for the AR0387 and AR0390 *TAC1B* allele swap strains was measured by qRT-PCR. In each graph, the median dCT value (calculated from the average *ACT1* gene CT value from three technical replicates subtracted from the average target gene CT value from three technical replicates) from three biological replicates of strain 1c or isolate AR0387, for panels A and B and panels C and D, respectively, was the comparator.

### *TAC1B*-mediated fluconazole resistance is driven predominantly by *CDR1* overexpression with a lesser contribution from *MDR1*

In order to further examine the contribution of *CDR1* and *MDR1* to fluconazole resistance due to *TAC1B* mutations, we disrupted these genes in resistant clinical isolate Kw2999 and strain 1c. *CDR1* disruption in Kw2999 resulted in a reduction in fluconazole MIC from 256 µg/mL to 4 µg/mL, and disruption in 1c resulted in a reduction in fluconazole MIC from 2 µg/mL to 0.25 µg/mL. *MDR1* disruption in isolate Kw2999 or strain 1c, however, had no effect on fluconazole MIC (**Figure 3**). Disruption of *CDR1* in strain 1cA640V resulted in a decrease in fluconazole MIC from 32 µg/mL to 0.5 µg/mL, disruption in 1c657V reduced the MIC from 32 µg/mL to 1 µg/mL, and disruption in 1cADdel reduced the MIC from 64 µg/mL to 0.5-1 µg/mL. As in Kw2999 and 1c, *MDR1* disruption had no effect on fluconazole MIC in any of these three strains (**Figure 3**).

**Figure 3.**
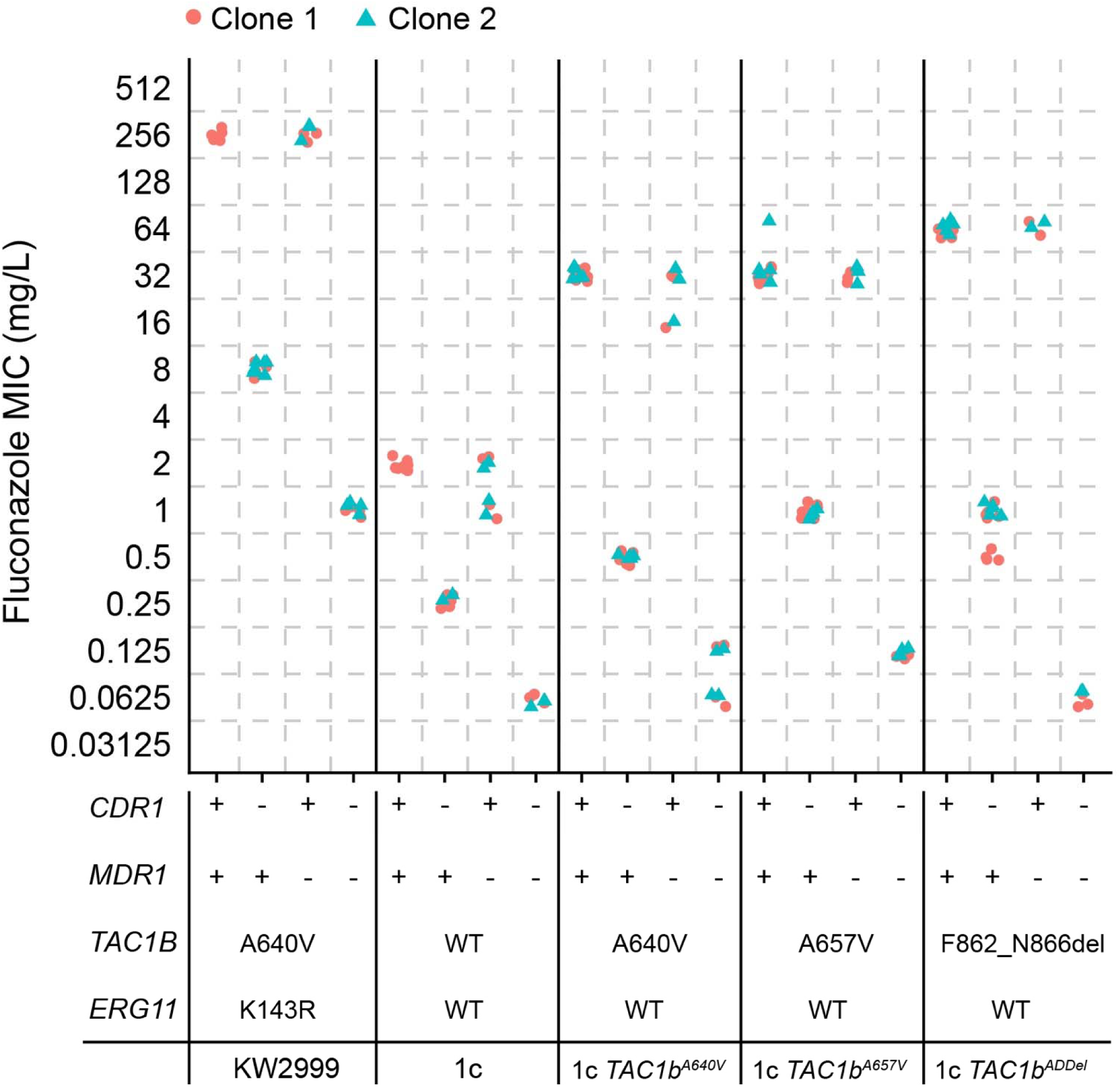
Fluconazole MIC were measured by broth microdilution in strains in which *CDR1* and *MDR1* were truncated by premature stop codon individually or in combination in Kw2999, 1c, or derivative backgrounds harboring indicated *TAC1B* mutations. Two independent clones were generated for each mutant combination. Each point represents a distinct MIC experimental replicate.

We also disrupted *MDR1* in the *CDR1*-disrupted derivatives of 1cA640V, 1cA657V, and 1cADdel and observed further reduction in fluconazole MICs for all strains tested. We observed a reduction from 0.5 µg/mL to 0.125 µg/mL in the 1cA640V background, 1 µg/mL to 0.125 µg/mL in the 1cA657V background, and 0.5-1 µg/mL to 0.0625 µg/mL in the 1cADdel background (**Figure 3**). In addition, disruption of *MDR1* in *CDR1*-disrupted mutants of Kw2999 and 1c resulted in a further MIC decrease (from 8 µg/mL to 1 µg/mL and from 0.25 µg/mL to 0.0625 µg/mL, respectively). Similar trends for all strains were observed for itraconazole, posaconazole, voriconazole, and isavuconazole (**Supplementary Figure S1**). These data indicate that the majority of fluconazole resistance conferred by mutations in *TAC1B* are driven by overexpression of *CDR1*, whereas *MDR1* makes a slight contribution to fluconazole resistance in isolates carrying such mutations.

## Discussion

Our findings reveal similarities and differences between genes influenced by *TAC1B* in *C. auris* and *TAC1* in the somewhat distantly related species *C. albicans*. In *C. albicans* Tac1 regulates the genes encoding the ABC transporters Cdr1 and Cdr2, both of which contribute to fluconazole resistance through activating mutations in *TAC1* leading to upregulation of these genes(Coste et al., 2004). In *C. albicans* Mrr1 regulates expression of the gene encoding the MFS transporter Mdr1 which also contributes to fluconazole resistance(Morschhäuser et al., 2007). In *C. auris*, *TAC1A* and *TAC1B* have been identified as homologs of *TAC1* and are situated in tandem with 488 bp between the ORFs on chromosome 5 of the *C. auris* genome(Mayr, Ramírez-Zavala, Krüger, & Morschhäuser, 2020). Only mutations in *TAC1B* have been implicated in fluconazole resistance, and such mutations have been widely identified across resistant clinical isolates in *C. auris* Clades I and IV and to a more limited extent in Clades II and III (Rybak et al., 2020).

We have previously shown that the A640V substitution in *TAC1B* leads to an increase in fluconazole MIC when introduced into a susceptible isolate and a reduction in MIC when the sequence is corrected to the wild-type sequence in a resistant isolate carrying this mutation(Rybak et al., 2020). In the present study we have confirmed the contribution of the A640V substitution and have shown that A657V and F862_N866del, have similar effects on fluconazole susceptibility. While ABC transporter gene *CDR1* overexpression in response to *TAC1B* mutations has not been previously established, the finding that this gene was upregulated among strains engineered to express these *TAC1B* mutations was not surprising given the relationship between *TAC1* and *CDR1* in *C. albicans* and other *Candida* species.

The overexpression of *MDR1* in strains carrying *TAC1B* mutations was unexpected and is reminiscent of the regulation of both *CDR1* and *MDR1* by Mrr1 in *C. lusitaniae* and both *CDR1B* and *MDR1B* by Mrr1 in *C. parapsilosis*(Demers et al., 2018; Doorley et al., 2022). Similar to *C. albicans*, to date, upregulation of *MDR1* in *C. auris* has only been associated with a mutation in *MRR1* present in all isolates of Clade III leading to a N647T substitution. Introduction of this mutation into a susceptible Clade IV isolate resulted in a nearly 80-fold increase in *MDR1* expression accompanied by an increase in fluconazole MIC from 4 µg/mL to 16 µg/mL and a similar fold increase in voriconazole MIC(Li et al., 2022). Deletion of the mutant *MRR1* in a clade III isolate resulted in a reduction in MIC from 512 µg/mL to 256 µg/mL and a similar fold reduction in voriconazole MIC. A similar effect was observed upon *MRR1* deletion in a different clade III isolate(Mayr et al., 2020). We observed at most an 11.7-fold increase in expression of *MDR1* in response to mutations in *TAC1B*. Given that disruption of *MDR1* alone in strains carrying *TAC1B* mutations had little to no effect on fluconazole MIC, and only a modest effect when disrupted in combination with *CDR1*, this comparatively modest level of *MDR1* overexpression appears not to be sufficient to have a substantive effect on fluconazole resistance.

Our work establishes the role of *CDR1* regulation through *TAC1B* mutations in fluconazole resistance in *C. auris* and clarifies the role of both *CDR1* and *MDR1* in isolates carrying such mutations. A more complete understanding of the repertoire of *TAC1B* and other mutations that contribute to fluconazole resistance in *C. auris* may one day lead to genetic tools to more rapidly and accurately predict patient responses to guide selection of antifungal therapy. Moreover, understanding how these mutations lead to *CDR1* upregulation and fluconazole resistance could point to therapeutic strategies for impeding expression of this efflux pump, thereby enhancing the activity and reclaiming the utility of fluconazole against *C. auris*.

## Supporting information

Online Supplement to Mutations in TAC1B

## Acknowledgements

This work was supported by NIH NIAID grant R01 AI169066 awarded to P.D.R. and C.A.C, NIAID grant U19AI110818 to the Broad Institute (C.A.C.), NIH NIAID grant T32 AI106700 (D.J.S.), and in part by the National Cancer Institute of the National Institutes of Health under Award Number P30 CA021765 awarded to the Hartwell Center at St. Jude Children’s Research Hospital.

