## Supplementary material for "Mutations in *TAC1B* drive *CDR1* and *MDR1* expression and azole resistance in *C. auris*": Online Supplement to Mutations in TAC1B

**Supplementary Methods.**

**Strain construction.** All oligonucleotides referred to below are listed by usage category in Supplementary Table S2. Strains were constructed as follows:

a. *Repair template design and preparation*. Forward and reverse primers for *TAC1B*^A640^ (wildtype), *TAC1B*^A640V^, *ERG11*^K143^ (wildtype), and *ERG11*^K143R^ manipulation, corresponding to the sequence surrounding the manipulation site with a silent mutation to destroy the PAM site and either containing the sequence resulting in the wildtype or mutant amino acid at the same position, were designed with a 20 bp overlap. Repair templates were prepared with these primers by primer extension (one cycle of 94^o^C, 5 min, 35 cycles of 94^o^C, 30s/42^o^C, 45s/72^o^C, 30s, and one cycle of 72^o^C for 5 min) in three 100 µl reactions containing 3 nmol each primer in 1x Phusion Green master mix (Invitrogen). The repair templates used to introduce the *TAC1B*^A657V^ or *TAC1B*^F862_N866del^ mutations to the wildtype sequence corresponded to 362 bp upstream and 387 bp downstream of the manipulation site for the A657V substitution, and 381 bp upstream and 366 bp downstream for the F862_N866del in-frame deletion, were synthesized as gBlocks by Integrated DNA Technologies (Coralville, IA). The repair template used for disruption of the *MDR1* ORF was synthesized as a gBlock corresponding to sequence 518 bp upstream of the manipulation site, a stop codon cassette consisting of 5’-TAGCTAGCTAG-3’ replacing the original sequence 7 bp downstream of the start codon, and sequence 521 bp downstream of the manipulation site. Similarly, the repair template used for disruption of the *CDR1* ORF was synthesized as a gBlock corresponding to sequence 490 bp upstream of the manipulation site, the stop codon cassette replacing the original sequence 88 bp downstream of the start codon, and sequence 488 bp downstream of the manipulation site. The gBlock repair templates were amplified using standard PCR conditions using Phusion Green master mix (Invitrogen/Thermo Fisher Scientific; Waltham, MA). PCR products were purified using a Qiagen PCR Purification kit per manufacturer’s instructions and eluted in 25 µl nuclease-free water (QIAGEN; Hilden, Germany).
b. *C*. *auris transformations*. Forty milliliters of an overnight culture initiated by inoculating YPD with a single colony of *C. auris* (final OD_600_=1.2-1.8) was used to prepare electrocompetent cells as described previously[14]. One microgram of guide-specific plasmid and 5 µg repair template were combined with 40 µl electrocompetent cells in a 0.2 cm BioRad Gene Pulser cuvette, and *C. albicans* programmed Gene Pulser settings were used to electroporate the cells (Bio-Rad Laboratories; Hercules, CA). Cells were allowed to recover in 0.5 ml 1M sorbitol + 1 ml YPD for 5 hrs at 35^o^C then spread on YPD + 200 µg/mL nourseothricin agar plates.
c. *Screening for positive transformants*. Initial screening of colonies from transformation plates was performed by colony PCR in 50 µl reactions containing 1x Phire Plant master mix (Invitrogen/Thermo Fisher Scientific) and using primers that bind sequences upstream and downstream of the repair template sequence. Purified PCR products were subjected to Sanger sequencing with locus-specific primers to detect the desired nucleotide change(s). Positive transformants were cultured in 5 ml YPD at 35^o^C for up to three passages of up to 72 hrs each to allow the cells to be cured of the pJMR19 plasmid. An aliquot of the culture was diluted 10^4^ and plated on YPD agar plates. Resulting colonies were replica plated on YPD agar and YPD + 200 µg/mL nourseothricin agar plates to screen for nourseothricin-sensitive colonies. Final confirmation of the entire sequence of the target gene was confirmed by amplifying the entire ORF plus ~200 bp of flanking sequence with locus-specific primers from genomic DNA isolated using MasterPure Yeast DNA Purification kit (LGC Biosearch Technologies; Middlesex, UK) in 50 µl reactions containing 1x Phusion Green master mix and subjecting the purified PCR products to Sanger sequencing.

**Supplementary Table S1. List of *C. auris* Clinical Isolates and Derivative Strains**

| **Isolate/Strain Name** | **Clone Designation** | **Relevant Genotype** | **Parent Strain** |
| --- | --- | --- | --- |
| Kw2999^†^ | N/A | *TAC1B*^A640V^/*ERG11*^K143R^ | Clinical isolate |
| 1c | N/A | *TAC1B*^WT^/*ERG11*^WT^ | Kw2999 |
| 1cA640V-1 | 1 | *TAC1B*^A640V^/*ERG11*^WT^ | 1c |
| 1cA640V-3 | 2 | *TAC1B*^A640V^/*ERG11*^WT^ | 1c |
| 1cA657V-7B1 | 1 | *TAC1B*^A657V^/*ERG11*^WT^ | 1c |
| 1cA657V-10A1 | 2 | *TAC1B*^A657V^/*ERG11*^WT^ | 1c |
| 1cADdel-10A | 1 | *TAC1B*^F862_N866del^/*ERG11*^WT^ | 1c |
| 1cADdel-4A | 2 | *TAC1B*^F862_N866del^/*ERG11*^WT^ | 1c |
| 2999*cdr1*-16A | 1 | *TAC1B*^A640V^/*ERG11*^K143R^/*CDR1*^S31*^ | Kw2999 |
| 2999*cdr1*-18A | 2 | *TAC1B*^A640V^/*ERG11*^K143R^/*CDR1*^S31*^ | Kw2999 |
| 1c-*cdr1*-6A | 1 | *TAC1B*^WT^/*ERG11*^WT^/*CDR1*^S31*^ | 1c |
| 1c-*cdr1*-8A | 2 | *TAC1B*^WT^/*ERG11*^WT^/*CDR1*^S31*^ | 1c |
| 1cA640V*cdr1*-4B | 1 | *TAC1B*^A640V^/*ERG11*^WT^/*CDR1*^S31*^ | 1cA640V-1 |
| 1cA640V*cdr1*-3A | 2 | *TAC1B*^A640V^/*ERG11*^WT^/*CDR1*^S31*^ | 1cA640V-1 |
| 1cA657V*cdr1*-22A | 1 | *TAC1B*^A657V^/*ERG11*^WT^/*CDR1*^S31*^ | 1cA657V-23A |
| 1cA657V*cdr1*-23A | 2 | *TAC1B*^A657V^/*ERG11*^WT^/*CDR1*^S31*^ | 1cA657V-23A |
| 1cADdel*cdr1*-8A | 1 | *TAC1B*^F862_N866del^/*ERG11*^WT^/*CDR1*^S31*^ | 1cADdel-6A |
| 1cADdel*cdr1*-6A | 2 | *TAC1B*^F862_N866del^/*ERG11*^WT^/*CDR1*^S31*^ | 1cADdel-6A |
| 2999*mdr1*-12A | 1 | *TAC1B*^A640V^/*ERG11*^K143R^/ *MDR1*^S5*^ | Kw2999 |
| 2999*mdr1*-17A | 2 | *TAC1B*^A640V^/*ERG11*^K143R^/ *MDR1*^S5*^ | Kw2999 |
| 1c-*mdr1*-8A | 1 | *TAC1B*^WT^ /*ERG11*^WT^/*MDR1*^S5*^ | 1c |
| 1c-*mdr1*-17A | 2 | *TAC1B*^WT^ /*ERG11*^WT^/*MDR1*^S5*^ | 1c |
| 1cA640V*mdr1*-8A | 1 | *TAC1B*^A640V^ /*ERG11*^WT^/ *MDR1*^S5*^ | 1cA640V-1 |
| 1cA640V*mdr1*-9A | 2 | *TAC1B*^A640V^ /*ERG11*^WT^/ *MDR1*^S5*^ | 1cA640V-1 |
| 1cA657V*mdr1*-7A | 1 | *TAC1B*^A657V^ /*ERG11*^WT^/*MDR1*^S5*^ | 1cA657V-23A |
| 1cA657V*mdr1*-11A | 2 | *TAC1B*^A657V^ /*ERG11*^WT^/*MDR1*^S5*^ | 1cA657V-23A |
| 1cADdel-*mdr1*-9A | 1 | *TAC1B*^F862_N866del^ /*ERG11*^WT^/*MDR1*^S5*^ | 1cADdel-6A |
| 1cADdel-*mdr1*-12A | 2 | *TAC1B*^F862_N866del^ /*ERG11*^WT^/*MDR1*^S5*^ | 1cADdel-6A |
| 2999*cdr1-mdr1*-7A | 1 | *TAC1B*^A640V^/*ERG11*^K143R^/*CDR1*^S31*^/*MDR1*^S5*^ | 2999*cdr1*-16A |
| 2999*cdr1-mdr1*-10A | 2 | *TAC1B*^A640V^/*ERG11*^K143R^/*CDR1*^S31*^/*MDR1*^S5*^ | 2999*cdr1*-16A |
| 1c-*cdr1-mdr1*-12A | 1 | *TAC1B*^WT^/*ERG11*^WT^/*CDR1*^S31*^/*MDR1*^S5*^ | 1c-*cdr1*-6A |
| 1c-*cdr1-mdr1*-38A | 2 | *TAC1B*^WT^/*ERG11*^WT^/*CDR1*^S31*^/*MDR1*^S5*^ | 1c-*cdr1*-6A |
| 1cA640V*cdr1-mdr1*-7A | 1 | *TAC1B*^A640V^/*ERG11*^WT^/*CDR1*^S31*^/*MDR1*^S5*^ | 1cA640V*cdr1*-3A |
| 1cA640V*cdr1-mdr1*-11A | 2 | *TAC1B*^A640V^/*ERG11*^WT^/*CDR1*^S31*^/*MDR1*^S5*^ | 1cA640V*cdr1*-3A |
| 1cA657V*cdr1-mdr1*-1A | 1 | *TAC1B*^A657V^/*ERG11*^WT^/*CDR1*^S31*^/*MDR1*^S5*^ | 1cA657V*cdr1*-22A |
| 1cA657V*cdr1-mdr1*-16A | 2 | *TAC1B*^A657V^/*ERG11*^WT^/*CDR1*^S31*^/*MDR1*^S5*^ | 1cA657V*cdr1*-22A |
| 1cADdel-*cdr1-mdr1*-3A | 1 | *TAC1B*^F862_N866del^ /*ERG11*^WT^/ *CDR1*^S31*^/*MDR1*^S5*^ | 1cADdel*cdr1*-8A |
| 1cADdel-*cdr1-mdr1*-14B | 2 | *TAC1B*^F862_N866del^ /*ERG11*^WT^/ *CDR1*^S31*^/*MDR1*^S5*^ | 1cADdel*cdr1*-8A |
| AR0387 | N/A | *TAC1B*^WT^/*ERG11*^WT^ | Clinical isolate |
| AR0387_*TAC1B*^A640V‡^ | N/A | *TAC1B*^A640V^/*ERG11*^WT^ | AR0387 |
| AR0390 | N/A | *TAC1B*^A640V^/*ERG11*^K143R^ | Clinical isolate |
| AR0390_*TAC1B*^WT‡^ | N/A | *TAC1B*^WT^/*ERG11*^K143R^ | AR0390 |

^†^ This clinical isolate was in the collection of isolates described in Ahmad et al. [28]

^‡^ These strains were previously described in Rybak et al. [15]

**Supplementary Table S2. Oligonucleotides used in this study**

| **Oligonucleotide Name** | **Oligonucleotide sequence (5’->3’)^†^** |
| --- | --- |
| ***sgDNA sequences for cloning*** | |
| TAC1B^A640WT^ TOP | ccaTCTCGTTCTTCGCCATGAAC |
| TAC1B^A640WT^ BOTTOM | aacGTTCATGGCGAAGAACGAGA |
| TAC1B^A640V^ TOP | ccaTCTCGTTCTTCGtCATGAAC |
| TAC1B^A640V^ BOTTOM | aacGTTCATGaCGAAGAACGAGA |
| TAC1B^A657V^ TOP | ccaGTTAGCAATCAAGTTCATCA |
| TAC1B^A657V^ BOTTOM | aacTGATGAACTTGATTGCTAAC |
| TAC1B^F862_N866del^ TOP | ccaCCTAATTTCTTCTTCGATAA |
| TAC1B^F862_N866del^ BOTTOM | aacTTATCGAAGAAGAAATTAGG |
| ERG11^K143(R)^ TOP | ccaCTGCTCCATCAACCTCGAGT |
| ERG11^K143(R)^ BOTTOM | aacACTCGAGGTTGATGGAGCAG |
| CDR1-STOP TOP | ccaCAACGGCAGTCTCAGTGAGG |
| CDR1-STOP BOTTOM | aacCCTCACTGAGACTGCCGTTG |
| MDR1-STOP TOP | ccaCAAAGTGTTCACATACCCCG |
| MDR1-STOP BOTTOM | aacCGGGGTATGTGAACACTTTG |
| Guide Check-F | GGGTGTCGGTTGGGTTGTG |
| ***Repair Templates (RT) and RT Amplification Primers*** | |
| ERG11^K143WT^ RT-F | GGGCACGTACCTCTGGAAAGCTTCTTTCGTCAAGGCAGTCTTAGCAAATTTCTTCTGCTCCATCAACCTCGAGTTtGGAC |
| ERG11^K143UNIV^ RT-R | CCGAGGCTGCTTATTCCCACTTGACCACTCCAGTTTTCGGGAAAGGTGTCATTTACGACTGTCCaAACTCGAGGTTGATG |
| ERG11^K143R^ RT-F | GGGCACGTACCTCTGGAAAGCTTCTTTCGTCAAGGCAGTCTTAGCAAATTTCcTCTGCTCCATCAACCTCGAGTTtGGAC |
| TAC1B^A640WT^ RT-F | CTTGACAGCGCAAGAACTATACTTCATCTTATTCGCGGCATCAATAGAAAAAGTGTCTCGTTCTTtGCtATGAAtTGGAT |
| TAC1B^A640WT^ RT-R | CGCTATCATCAGAATAATTGAGGCAGTTAGCAATCAAGTTCATCATGGCGAAAAAAGGATAAGTAATTATCCAaTTCATaGCaAAGAACGAGAC |
| TAC1B^A640V^ RT-F | CTTGACAGCGCAAGAACTATACTTCATCTTATTCGCGGCATCAATAGAAAAAGTGTCTCGTTCTTtGttATGAAtTGGAT |
| TAC1B^A640V^ RT-R | CGCTATCATCAGAATAATTGAGGCAGTTAGCAATCAAGTTCATCATGGCGAAAAAAGGATAAGTAATTATCCAaTTCATaaCaAAGAACGAGAC |
| TAC1B^A657V^ gBlock | TAAGAATATATCGAATGAGTTGAAGGAAGAGTTTCGTCCTCGCTTTTACTTCGAGCCTGAATTCGCGTTAATGTTGGCGAGATTTTCCGGAGATGAAAGGATGAGCAGCGTCAATGATGGTATTCTTTCGTTCCAGCTTGCCTACTTCTTCCAGCTCATGACCATCAATAAGGTTCCTTCGCAGCTCGACCCCAATCAAGGTAATACCCCTCCTTATGAAAACTCACAGTACAGAAAACTTGCCCTTGACAGCGCAAGAACTATACTTCATCTTATTCGCGGCATCAATAGAAAAAGTGTCTCGTTCTTtGCtATGAAtTGGATAATTACTTATCCTTTTTTCGCtATGATGAACTTGATTGtTAACTGCCTCAATTATTCTGATGATAGCGAAGGCGTTACCGACTTGAACTTGCTTATTGATTTATCAATGAACTTCTTTAATCACTACACTGATTTGGCTAAGAAACCCTCAACCGGTGCATTCTACTTACGTTTGCATTTATTTGGCATTATTATTCGTATTGCTCTTCGTATCACTGTGAAAGTATATGAAGAAAATAACAATGTTGACATATTGGGCAGCAACCCTCAGCTTAAGTCGCATTTAGAACAAGTTGAAAAGGAGTTTCCTCAATTTTACACAGAAGTGAGCGCTCCTTCCGATTTGTTGAGTCTCTTGACTTGCATGCATCCGTACACAGATATAAATCTGCAAAGTGACGACAGGAAATTCACGCCCAATGGTTC |
| TAC1B^A657V^ RT_Amp-F (identical to TAC1B SCN_Seq-F) | CCTCGCTTTTACTTCGAGCC |
| TAC1B^A657V^ RT_Amp-R | GGGCGTGAATTTCCTGTCG |
| TAC1B^F862_N866del^ gBlock | TCGCATTTAGAACAAGTTGAAAAGGAGTTTCCTCAATTTTACACAGAAGTGAGCGCTCCTTCCGATTTGTTGAGTCTCTTGACTTGCATGCATCCGTACACAGATATAAATCTGCAAAGTGACGACAGGAAATTCACGCCCAATGGTTCGCTTGATACATCCTCGAGCGTATCAAACCAAGACATTGCTTCGCTAACTTTTCAACACGCGAGCGGCATGGAGGCACCTTCACCCAAAAAGAATGACCCTACACTTTCCAACATACTTCATCCTGTTGACTTTGCATCTGCTAGAGATTCTGAAAGAAAAGACTTCAATATCGACGACGACCTATTGCTCGCCGCGATAAACCAAGATTTTCTGGCACTTCCTAATGGGCTTTAACTTTGTAAATAGTATGCTTACCACGATTTTAACGATGATGATATACTTTTATACTAATTCCACATATGCTAACGAGTACGAATTACGCCAGCAGCAATCAACATGTTGACTTTCTTTGCCACCTCTGGGTTCTTCATGTGCTCTTGCAATGCCGCTGGATTATCTCTGGCTTGACCCAATATACCCTGCATGACAGGGTCTTGCAAAATTTCAACTATTTCTGGGTCCTTAGACACTCTCTCCATCGTTTGTTCCGGAGTTTCGCCCTCGATTGCTGCAAAACGCTGCGACATTGCTCTGTGCATCAACTGGTTGATTTCGTTGGCGTTCTTTCCACCGTTCAACTCTTGATCCTTTGCAAGAGCG |
| TAC1B^F862_N866del^ RT_Amp-F | GAAGTGAGCGCTCCTTCCG |
| TAC1B^F862_N866del^ RT_Amp-R | CGCCAACGAAATCAACCAG |
| CDR1-STOP gBlock | GGCAAATTGTCCCGCTGGAAGTGGAGACTCCAGTTGTTGCCCACAGGTTATGAAGCTGGTGGTTATTTTTGTTGTATGTGGTTAGGGCTGAGAAGGGCGGTTTTTTTGTGGCATTTTGTCCCCAGTGATTAAATGTGCACCGCCAGGCTCCCGGTGGGAGTGGCCAGAGCCGGAACTCGCACCTTCCGAAGTTCCACCGCCAGCCGTCTCCAACGTGCGCAACTCTCGCACAATACGCTATTTGCCGCCAGAATTCCCTATCTCATGCCTGCTGCTCCTGCAGCGTAGCTGGGGCTGCTTCTACCTCTCCCTCTGCGACATTGTTTGGCATTTCTTTTCGCCTGCCAATATATATATCGCATGTAGTCCACTTTAAAGGCAAATCTTCACTTTTCCATCTCCATGTCCGAGAAACCTTTTGTCGACGCTCCTCCACCCGAGGATGGCGTTGCTCACCAAGTGCTGCCCCATGACAACGGCAGTCTCAGTGctagctagctagAGGAGGCCAATTCCATCAATGAGTATACTGGTTTTGGTGCTCATCAGGAAGGTGAAATTAGAGAGTTGGCCAGAACCTTCACCAACATGTCCCATGACTCCGGCCACGACTTATCCAAAACAAACACATCCCAGGATTTGCTCAAGTACTTGTCCCACATGTCTGAGGTGCCTGGCGTAGAGCCCTTTGACCCAGAGCAGATCAGCGAGCAGTTGAACCCAGACTCGCCCAACTTCAATGCGAAGTTTTGGGTGAAAAATATGCGTAAGTTGTTCGATTCCAACCCTGACTACTATAAGCCTTCAAAGTTGGGACTTGCGTACCGTAATTTGAGAGCCTACGGTGTGGCTGCAGACTCAGACTACCAGCCAACCGTCAGTAACGGGTTGTGGAAAATGGCGGTGGATTACTGGCACGATATGAGAAAAATCGACGAGAGCCGTTGTTTTGACATCTTAAAGACCATGGACGGGTACTTCAAGCCCGGT |
| CDR1 RT_Amp-F | CCGCTGGAAGTGGAGACTCC |
| CDR1 RT_Amp-R | GCTTGAAGTACCCGTCCATGG |
| MDR1-STOP gBlock | GGTTCACGTAAGCGACCCTTCGTACGGTCCTGTCAATATGCAGCACGTACCCGTCGTTCGGCGCCTACCATTTTAACCCGGAGATGCGACAAGGGCACAAAAAGCGACTAGTCATTGCGCAAGCAAACGTGTGAAAGTGACTCTAAAAGACTGATTTAGGGAGACCTGGAAGGAGTTCGAAAAGCTCATGAAACTAAAAAATAAAATGAATAATTAAAAAAAAAAAAAAAAAGCAATGCCTCGTACTCCCGTCAAACTCACGTAAGCCTTTTCGTCTGCACCCAAACAAAGACAATCTAACACTTTCGTACATTTTCTGCGCGGCTTAACTCAAATGTGGCGGCTGCAAAAGGTATGCCATTTTTGCTCTCATTACTCATCAAAAAAATAGTGATAAGGTACACTGAGACATATAAATAGGCTGTAACTTCTCTGATTCTTTCGTCTTTACCAAATTTCAACGCATCTTCAATCTCCACATGTTCCTagcTAgcTagGTCAGAGAGAGCTTCTTCGGCAGGTCTCTATATCACTTATCTGGACGCAAAGTGTTCACATACCCCGAGGAATCACCCGACTATGTGATTCCCGCAAAGTACTTGGGCAAAGACGAAGCAGGCATTGAATCGGATGTTCAGGAGAAAGCTGGCGCTTCAGACACCCCCGTTGATCTGGACTCTTCGTCCCAGTCGACCAAGACCAACCACATTCTTGTGGACTGGGAGGGAGAAGACGATCCAGAAAATCCATACAATTGGCCATTGAAATACAAGATTATCTTCATTGCCCAGATCATGATTTTGACTGCATTTGTGTATATGGCTTCTGCCATTTATACCCCAGGTATTGAGGAGATTATGAAAGATATGGGCGTGGGCCAGGTGGTGGCGACACTCCCCTTGACGCTTTTTGTGTTCGGATACGGTATCGGGCCCATGGTGTTTTCGCCGCTTTCGGAAAATGCCAGGTTTGGCAGAACGTCCATCTACATCATTACCTTATTTATCTTCTTTATCTTGCAGATCCCCACGGCTCTCGT |
| MDR1 RT_Amp-F | GCGACCCTTCGTACGGTC |
| MDR1 RT_Amp-R | CCGTGGGGATCTGCAAG |
| ***Screening and Sequencing Primers*** | |
| TAC1B SCN_Amp-F | CAAGTTGTCATCTCGGCCATC |
| TAC1B SCN_Amp-R | CAACATGTTGATTGCTGCTGGC |
| TAC1B SCN_Seq-F | CCTCGCTTTTACTTCGAGCC |
| TAC1B SeqF1 | CCTCCTGCGCCTTCCAC |
| TAC1B SeqF2 | CAACAACGCTTCCAATGAGCC |
| TAC1B SeqF3 | GAGTTACCGATCTGCGCCCC |
| TAC1B SeqF4 | CACCTCTCTTAACGGCGAAC |
| TAC1B SeqF5 | CATATTGGGCAGCAACCCTC |
| CDR1 5’ SCN_Amp-F | GAGGCAGCCCACTGTATGG |
| CDR1 ORF SCN_Amp-R | GTGGCATCCAAGATGGCCG |
| CDR1-STOP SCN_Seq | GCTTGAAGTACCCGTCCATGG |
| ERG11 SCN_Amp-F | CTCTTGGACCAAAAACAGCCAGC |
| ERG11 SCN_Amp-R | GGCTGGAGCTGGTTTGGTG |
| ERG11 SCN_Seq-F | GCAGAGAGAAATACGGCGATGTG |
| ERG11 SeqF1 | CCATCGTATAGTGGTCCTCC |
| ERG11 SeqF2 | GGGACTTGATCGACTCGC |
| MDR1-STOP SCN_Amp-F | CGGCCGTATATGTGGGCTC |
| MDR1-STOP SCN_Amp-R | CCCGTGAGAGCTCTGAGTC |
| MDR1dis SCN_SeqF | GGGCGATTCTCCGGTGG |
| ***qRT-PCR Primers*** | |
| ACT1 qRTPCR-F | GAAGGAGATCACTGCTTTAGCC |
| ACT1 qRTPCR-R | GAGCCACCAATCCACACAG |
| CDR1 qRTPCR-F | GAAATCTTGCACTTCCAGCCC |
| CDR1 qRTPCR-R | CATCAAGCAAGTAGCCACCG |
| MDR1 qRTPCR-F | GAAGTATGATGGCGGGTG |
| MDR1 qRTPCR-R | CCCAAGAGAGACGAGCCC |

†- lower-case nucleotides represent nucleotide changes from the original sequence

**Figure S1. Itraconazole, posaconazole, voriconazole, and isavuconazole MIC values from *TAC1B* mutant strains and their derivative *cdr1*-, *mdr1*-, and *cdr1/mdr1*-disruptant strains**

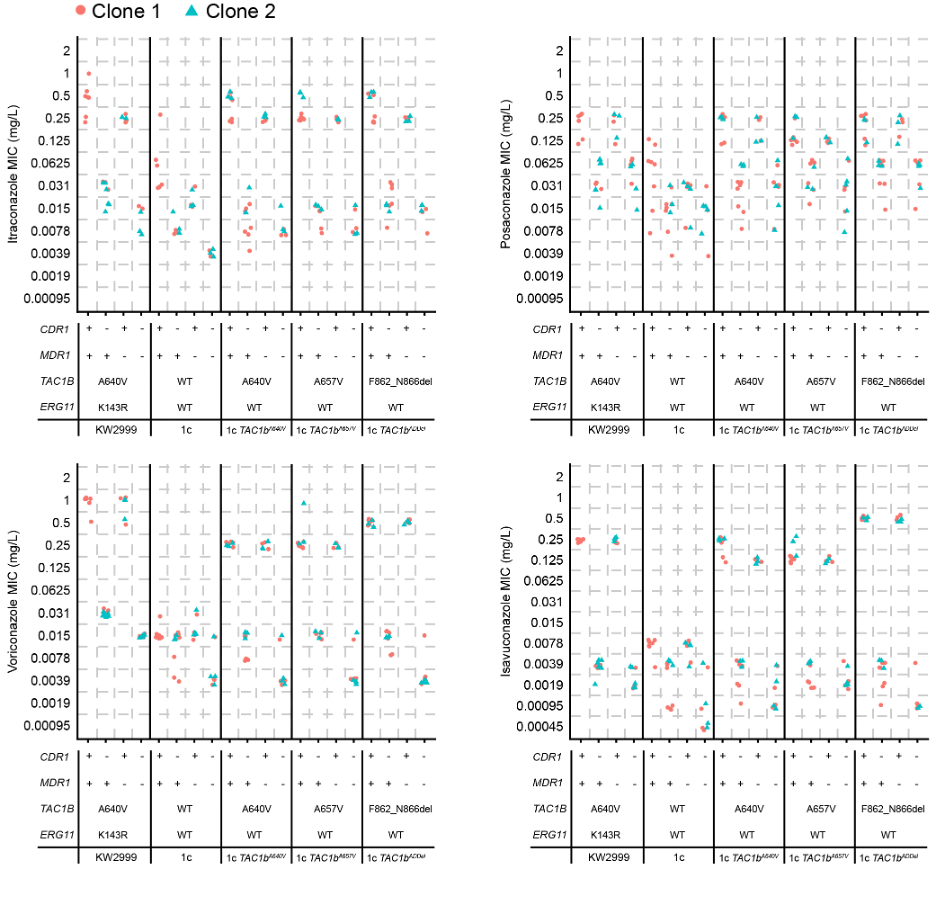

A.

B.

C.

D.

**Figure S1.** Itraconazole **(A)**, posaconazole **(B)**, voriconazole **(C)**, and isavuconazole **(D)** MIC were measured by broth microdilution in strains in which *CDR1* and *MDR1* were disrupted by premature stop codon individually or in combination in Kw2999, 1c, or derivative backgrounds harboring indicated *TAC1B* mutations. Two independent clones were generated for each mutant combination. Each point represents a distinct MIC experimental replicate.
